# Beyond climatic drought indices : an hydraulic approach to quantifying forest water stress

**DOI:** 10.64898/2026.07.13.738371

**Authors:** Hervé Cochard, Maxime Cailleret, Médéric Aubry, Phillipe Balandier, Arsène Druel, Marylou Mantova, Ludovic Martin, Thomas Opitz, Léo Place, Nicolas Martin-StPaul

## Abstract

The article introduces a new Forest Stress Index (ISF) based on a plant hydraulic modelling approach rather than classical climatic drought indices. Unlike other index like scPDSI or SPEI, ISF is grounded in xylem embolism dynamics simulated with the mechanistic SurEau model. The goal is to better link climatic anomalies to tree physiological functioning and mortality risk. ISF is defined using a locally adapted ideotype characterized by an optimal P_50_ value under a reference hydraulic functioning threshold. Simulations are performed across Europe and France using multiple climate datasets. The index is robust to model parameterization choices and assumptions about plant functional traits. Results show strong spatial and temporal consistency and significant correlations with SPEI and scPDSI. However, ISF more strongly highlights extreme drought years and exhibits a more skewed distribution. Future projections under SSP5-8.5 indicate a widespread increase in hydraulic stress with strong regional contrasts. Overall, ISF provides a mechanistic and complementary drought indicator more directly linked to forest mortality processes.

## Introduction

Forests worldwide are currently facing a rapid increase in the frequency, intensity, and duration of droughts as a result of climate change. These events are causing growth reductions, widespread decline, and large-scale forest mortality episodes across many biomes. In this context, the quantitative characterization of forest water stress has become a major challenge for understanding decline mechanisms, predicting ecosystem vulnerability, and assessing their capacity to adapt to future climatic conditions. Drought is usually described using meteorological indices that characterize the magnitude and duration of precipitation deficits—sometimes combined with anomalies in atmospheric evaporative demand (AED)—relative to the long-term climatology. Numerous drought and water stress indices have therefore been developed over the past decades (Tramblay et al 2020). The first indices used in forest ecology were mainly climatic or meteorological. Among the most widely used are the Standardized Precipitation Index (SPI, (McKee et al., 1993), the Palmer Drought Severity Index (PDSI, Guttman et al., 1998)), and more recently the Standardized Precipitation Evapotranspiration Index (SPEI, VicenteSerrano et al., 2010)), which explicitly incorporates atmospheric evaporative demand through potential evapotranspiration. The SPEI has notably become a benchmark indicator in global studies of forest decline because it simultaneously accounts for precipitation deficits and the increase in atmospheric evaporative demand associated with climate warming (Tramblay et al 2020 ; Bharghavi et al 2025).

However, these climatic approaches present fundamental limitations when it comes to representing the actual drought stress experienced by trees and the associated risk in forest ecosystems. Indeed, they mainly describe a meteorological or hydrological anomaly, without explicitly representing their impact on the functioning of trees. Thus, the same SPEI value or water deficit may have very different biological consequences depending on the species, forest stands, or biomes considered. A moderate drought in a humid temperate forest may cause severe damage, whereas much drier climatic conditions may remain close to the normal functioning range of Mediterranean or xerophytic ecosystems adapted to drought. This limitation arises from the fact that most conventional indices implicitly assume identical sensitivity thresholds among ecosystems, even though tree physiological traits vary strongly along climatic gradients.

Significant progress has been made in recent years in understanding the physiological processes involved in tree mortality under drought and heat stress. Current scientific consensus converges on the risk of hydraulic failure of the xylem water transport system by embolism as the key process underlying tree mortality (IPCC 2022). The work of Choat et al (2012) notably showed that many forest species operate with relatively narrow hydraulic safety margins regardless of the biome considered, suggesting a global convergence in the hydraulic vulnerability of forests. Thus, species living in the most arid environments generally exhibit more negative water potentials but also xylem tissues that are more resistant to embolism, thereby limiting the risk of hydraulic failure. Thus, the level of xylem embolism appears to be a mechanistic indicator of the water stress experienced by a tree in situ, directly linked to the risk of mortality induced by hot drought events. Building on these insights, the sequence of physiological impairments that develops during drought has been elucidated (Martin-StPaul et al. 2017; Choat et al. 2018) and subsequently incorporated into the mechanistic model SurEau (Cochard et al. 2021; Ruffault et al. 2022).

In this study, we propose a new forest stress index, termed ISF (Forest Stress Index), based on anomalies in rate of xylem embolism relative to a locally adapted hydraulic ideotype. The approach relies on defining, at each geographical location, an ideotype tree characterized by a combination of hydraulic traits optimized for local climatic conditions. This ideotype is defined such that it exhibits a mean embolism rate above a threshold value (typically 12%) over a historical reference climatic period representing optimal thriving conditions. Thus, unlike conventional indices based on absolute drought thresholds or hydraulic dysfunction, the ISF expresses water stress as a relative anomaly with respect to the expected functioning of a forest system theoretically adapted to the local reference climate. The recent development of the mechanistic model SurEau for predicting embolism risk now provides the opportunity to calculate this new water stress index.

## Materials and methods

In this study, we present the main principles of the ISF index. We then provide computation at the scale of Europe and, with finer spatial resolution, at the scale of metropolitan France. This calculation requires access to climatic data and an appropriate parameterization of the SurEau model.

### The Sureau model and its application in computing ISF

*SurEau* is a mechanistic model of tree water and hydraulic functioning based on the explicit representation of the physiological processes involved in water transport and gas exchange. It enables the simulation of the temporal dynamics of transpiration fluxes, stomatal conductance, water potentials, and tree water storage under a given soil and climatic environment. The model was specifically developed to predict the dynamics of xylem embolism and thereby quantify the risk of hydraulic failure that may ultimately lead to tree mortality during drought events. The *SurEau* model was described in detail by Cochard et al. (2020), Martin St-Paul et al (2017) and Ruffault et al 2022. For the present study, we used version 2026_05_03 of *SurEau*.*c*. In this version, we incorporated a model describing the dynamics of snowpack thickness as well as soil temperature, based on heat exchanges between the soil and the surrounding atmosphere. When the temperature of a soil layer falls below 0 °C, the water uptake capacity of the roots located in that layer is assumed to be completely inhibited. Consequently, the model predicts an increase in water stress and xylem embolism during winter when the soil is frozen while air temperature remains positive, thereby allowing residual foliar evapotranspiration.

### Climatic data

The SurEau model uses daily climatic input variables including temperature, humidity, radiation (PAR), precipitation, and wind speed. Several datasets were mobilized in this study to provide these climatic forcing variables, allowing us to assess the sensitivity of ISF estimates to the choice of climate data source.

1. The <u>SAFRAN</u> dataset from Météo-France is an atmospheric reanalysis that provides homogenized surface meteorological variables over France on a regular grid of approximately 8 × 8 km. Daily data are available from 1959 to present.
2. The <u>E-OBS</u> is a high-resolution gridded observational dataset produced by interpolating in-situ weather station measurements across Europe, providing daily values of surface climate variables on a regular grid of approximately 0.1° (∼10 km), with coverage starting in 1950 to the present. Data were retreived from the Copernicus Climate Data Store. It should be noted that wind speed values are only available from 1980 onwards in this dataset. Prior to this date, wind speed is assumed to be constant and set to a mean value of 2 m s^−1^.
3. The <u>ERA5-Land</u> is a high-resolution land surface reanalysis produced by the European Centre for Medium-Range Weather Forecasts, providing hourly meteorological and land surface variables at a spatial resolution of about 9 km, derived from a combination of physical modeling and data assimilation of reanalysis atmospheric fields, with global coverage from 1950 to present on Copernicus Climate Data Store.
4. For the calculation of the ISF under future climate projections, we used the climate model developed by the <u>EC-Earth3 CMIP6</u> consortium within the CMIP6 framework. This model provides physically based simulations of the coupled atmosphere–ocean–land climate system, driven by greenhouse gas concentration scenarios used in climate change projections. Daily climate data are available on a global grid at a spatial resolution of 0.5° for the period 1950– 2100. In this study, we selected the SSP5-8.5/r1i1p1f1 scenario dowloaded from the NASA/NCCS THREDDS Data Catalog.

Whenever necessary, climatic data units were converted in advance to match the format required by the SurEau model.

### Main principles and hypothesis underlying the ISF

The ISF is computed in two main steps (figure 1). First, the model is calibrated to define a plant ideotype under reference climatic conditions. Second, SurEau is run with this ideotype over a given climate time series to simulate the daily progression of xylem embolism.

**Figure 1:**
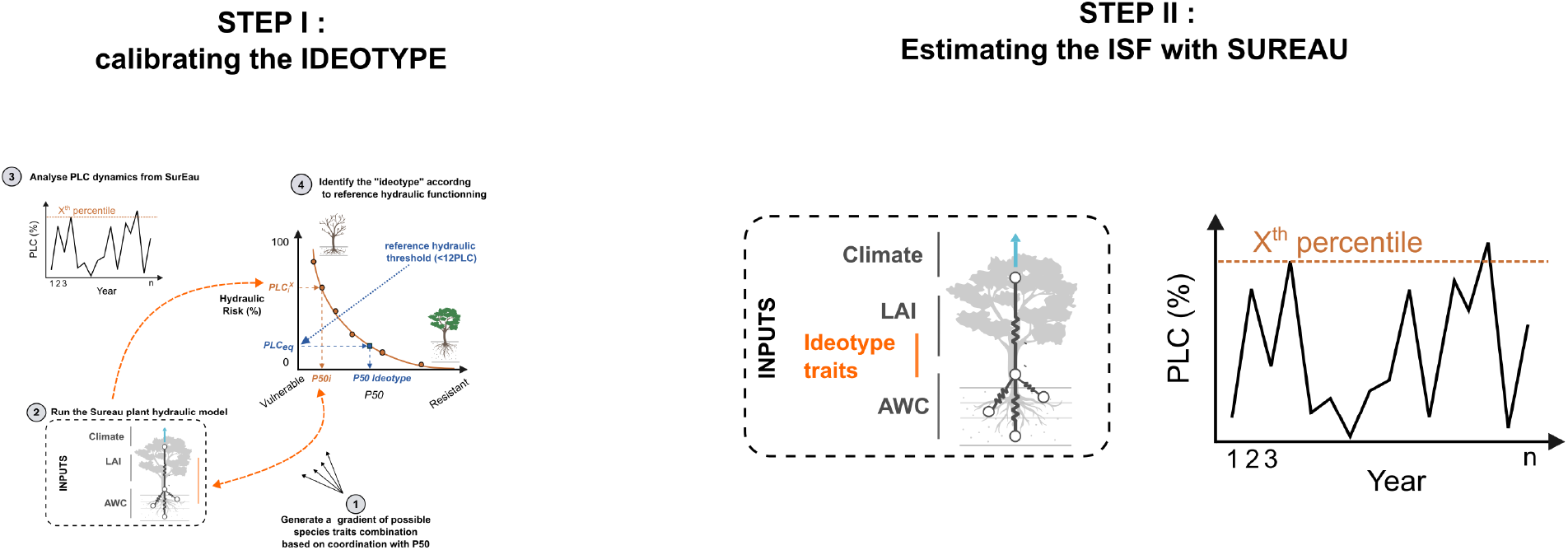
Workflow used to calculate the Forest Stress Index (ISF). First, the SurEau model is calibrated under reference climatic conditions to determine the hydraulic traits of a local plant ideotype (*P*_*50*__id) that maintains a mean annual percent loss of hydraulic conductivity (PLC) of 12%. Second, the calibrated ideotype is forced with the target climate time series to simulate the daily dynamics of xylem embolism. The annual ISF is then calculated as the anomaly in mean annual PLC relative to the reference period.

For most ISF calculations, we considered a standard mature evergreen tree 25 m tall and 30 cm in diameter (DBH). We also considered the case of a deciduous tree, as well as an evergreen shrub 7 m tall with a DBH of 7 cm. Following the hydroecological equilibrium hypothesis proposed by Eagelson (1982) and Druel et al (2026), we kept leaf area index (LAI = 4 m^2^ m^−2^) and total available water in the soil (TAW = 100 mm) constant across all grid points. However, we tested the impact of a ±20% variation in LAI and TAW on ISF.

The ISF calculation is based on identifying an idealized tree phenotype whose combination of hydraulic and water-use traits is optimal for given climatic conditions. Here, we defined this physiological optimum as the mean reference embolism level (PLC_ref = 12%) reached each year over a reference period. We also tested the impact of a ±20% variation in PLC_ref on ISF. Because the effects of these physiological traits on embolism can compensate for one another, there is an infinite number of trait combinations consistent with the optimality hypothesis we adopted. To circumvent this issue, we varied only xylem resistance to embolism (P_50_, MPa) and assumed empirical correlations with the other key traits. These relationships were established from available trait databases (e.g., Choat et al 2012, Martin St-Paul et al 2017).

Thus, we assumed that the slope of the vulnerability curve at P_50_ (slope, %), residual leaf conductance (g_res, mmol s^−1^ m^−2^), the leaf water potentials inducing 12% and 88% stomatal closure (P_gs_12_ and P_gs_88_, MPa), and the osmotic potential at full turgor (Π_0_, MPa) were related to P_50_ according to the following equations (supplemental figure S1):

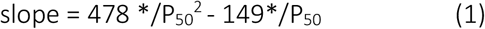

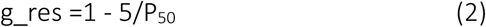

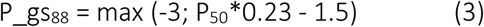

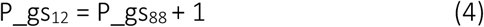

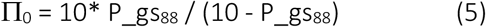

All other traits and model parameters were considered constant. As a consequence of this parameterization, the time to hydraulic failure (THF, days) increased as P50 decreased, regardless of air temperature (Figure 2). The sensitivity of THF to P_50_ was greater at higher P_50_ values.

**Figure 2:**
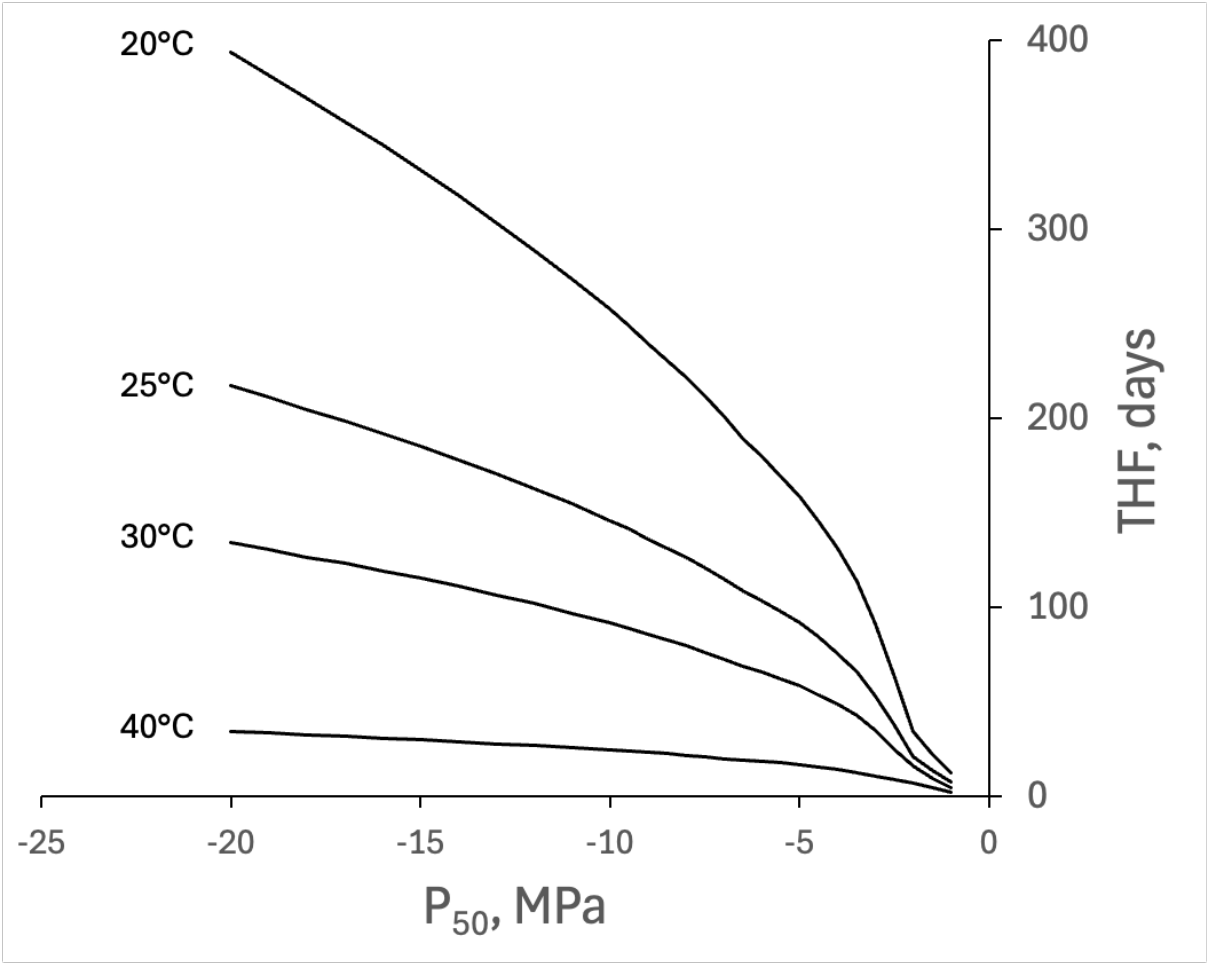
RelaLonship between the xylem vulnerability to embolism (P_50_) and Lme to hydraulic failure (THF) as predicted by the SurEau model. For each P_50_ value, key other model traits were adjusted according to their empirical covariaLon with P50 The relaLonships are shown for four air temperature condiLons.

### Computing the ISF

The ISF calculation first consists of identifying, for each grid cell, the P_50_ of the ideotype (P_50__id) that satisfies the constraint PLC_ref = 12%. To do so, we used a historical reference period (1960–1969) common to the different climate datasets. For the ERA5 dataset, we also considered a longer period (1950–1969), which had no significant effect on P_50__id or ISF. This reference period is assumed to be representative of a historical climatic baseline compatible with the climatic niche of the tree ideotype being identified. For each grid point, the model is first initialized with an initial P_50_ value (typically −5 MPa) and the associated traits. If, at a given grid point, the mean PLC over the reference period is higher than the target PLC_ref, then P_50_ is decreased ; otherwise, it is increased. This iterative procedure is repeated until convergence to PLC_ref is achieved, with a tolerance of ±0.5%, hence defining P_50__id. The ISF is defined as the annual embolism rate of ideotypes at each grid point for any climatic time series. By construction, ISF is equal to 12% over the reference climatic period, lower than 12% in wetter years, and may reach 100% under extremely dry conditions.

### Reference drought index

We compared ISF values with two reference drought indices : the Self-Calibrating Palmer Drought Severity Index (<u>scPDSI</u>; Wells et al., 2004) and the Standardized Precipitation–Evapotranspiration Index (<u>SPEI</u>; Vicente-Serrano et al., 2010). August scPDSI values for each year were extracted from Barichivich et al. (2025). August SPEI values, calculated over a six-month timescale, were extracted from Beguería et al. (2010). These indices were derived from climate datasets that differ from those used to calculate ISF ; however, differences between the climatic forcing datasets are generally small over France and are therefore unlikely to substantially affect the comparison.

## Results

### Robustness of the ISF index

In a first set of simulations, we assessed the robustness of the ISF with respect to the choices made in parameterizing the SurEau model. To this end, we selected two contrasting grid points, one located in central France (2.68°E, 46.53°N) and the other in central Spain (−3.32°E, 41.53°N). The ERA5_Land climate dataset was used for this analysis, with a reference period of 1950–1969. The corresponding P_50__id values were estimated at −3.97 MPa for France and −7.43 MPa for Spain. We then computed the ISF over the 2010–2025 period by varying several key model parameters.

A positive relationship was observed between TAW and P_50__id, with higher soil water storage associated with less negative P_50__id values (figure 3a, insert). Across the explored range of variation (±20%), the sensitivity of P_50__id to TAW was 0.30 %%^-1^ in France and 0.46 %%^-1^ in Spain. However, after correcting P50_id for TAW, the ISF becomes independent of TAW at both sites (Figure 3a).

**Figure 3:**
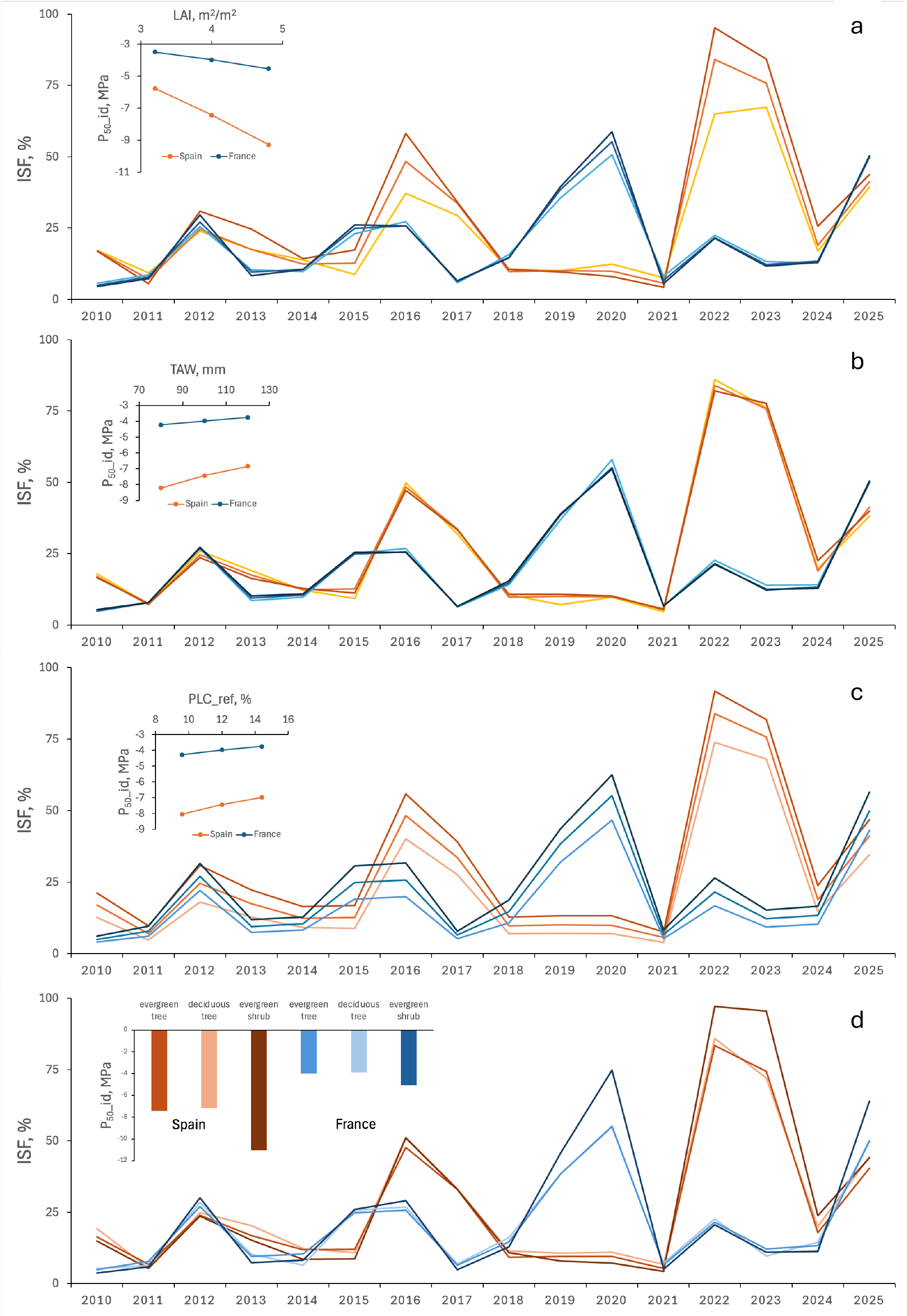
Sensitivity analysis of P50_id and ISF to key SurEau model parameterizations at two contrasting sites (central France : 2.68°E, 46.53°N; central Spain : −3.32°E, 41.53°N) using ERA5-Land climate data (1950–1969 reference period; ISF computed over 2010–2025). Panels show the effects of variations (±20%) in (a) total available soil water (TAW), (b) leaf area index (LAI), (c) reference embolism threshold (PLC_ref), and (d) plant life form (evergreen tree, deciduous tree, shrub) on P50_id (insets) and ISF. P50_id is strongly sensitive to model parameters, particularly LAI and TAW, whereas ISF remains largely robust, with only limited impact on interannual variability and ranking of extreme stress years.

Similarly, P_50__id responded strongly and negatively to variations in LAI (Figure 3b, inset), with increasing LAI leading to more negative P_50__id values. This response was more pronounced under arid conditions (sensitivity of −1.18 %%^-1^) than under temperate conditions (−0.66 %%^-1^). In contrast, the effect of LAI variations on ISF was strongly attenuated, although not entirely compensated for as observed with TAW (Figure 3b). Nevertheless, the ranking of the years with the highest ISF values remained globally unchanged. Changing the value of PLC_ref also affected P_50__id (figure 3c, insert), with higher PLC_ref values leading to slightly less negative P_50__id values (sensitivities of 0.36 %%^-1^ and 0.33 %%^-1^ for Spain and France, respectively). A 20% variation in PLC_ref induced a nearly proportional systematic change in ISF (18.4%) at both sites (figure 3c).

We further investigated the impact of ideotype life form (evergreen vs. deciduous ; tree vs. shrub) on ISF and P_50__id. Tree deciduousness increased P_50__id by only 3.9% in Spain and 2.1% in France, with only a marginal effect on ISF (Figure 3d). In contrast, considering a shrub rather than a tree as the life form drastically decreased P_50__id, with an even stronger effect in dry (−35.3%) than in temperate (−23.2%) habitats. Shrubs also exhibited higher ISF values ; however, the ranking of the years with the highest ISF values remained globally unchanged.

In a final set of simulations, we compared the three climate datasets. These datasets exhibited non-negligible differences in their climatic estimates, leading to substantial variations in the predicted ideotype P_50__id. In central France, for example, P50_id was estimated at −3.43 MPa using the Safran dataset, −3.60 MPa with E-Obs, and −3.97 MPa with ERA5-Land. Nevertheless, interannual variations in the ISF remained relatively consistent across datasets, with the years 1976, 2003, and 2019–2020 systematically emerging as periods of particularly high stress (Figure 4).

**Figure 4:**
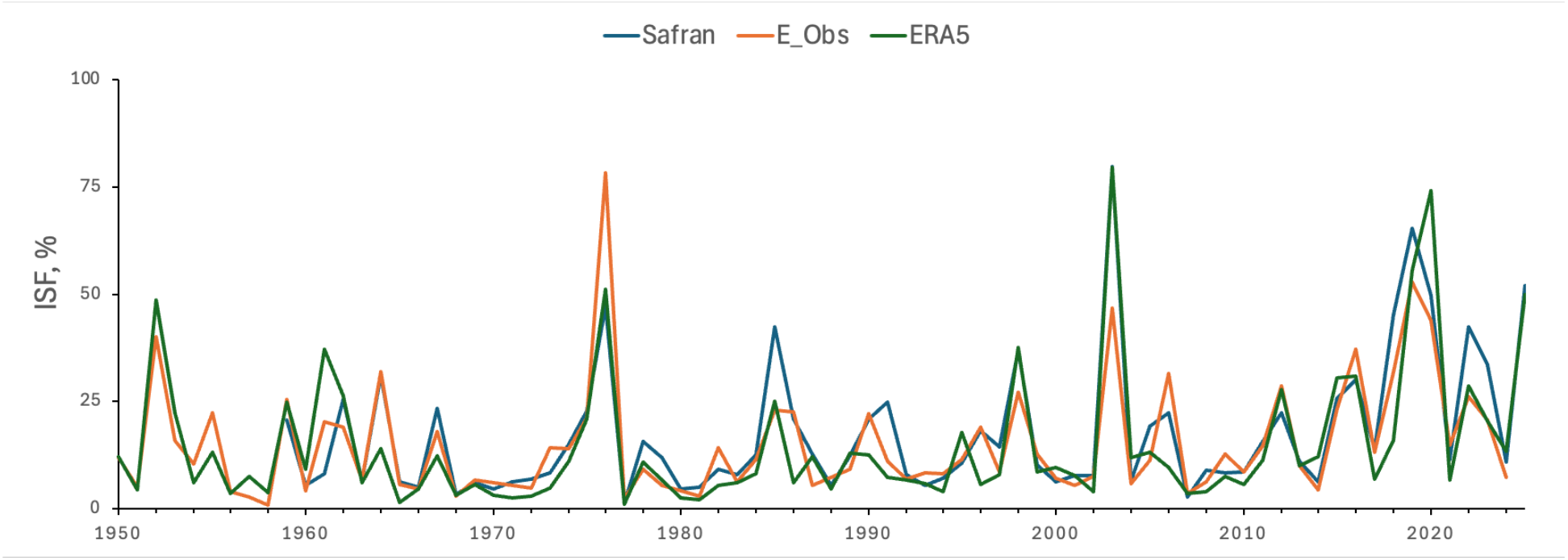
Comparison of the influence of different climate forcing datasets on P50_id and ISF estimates. Results are shown for central France using SAFRAN, E-OBS, and ERA5-Land climate datasets.

### Spatial variation in P_50__id across France and the Euro–Mediterranean region

P_50__id values were first computed on a grid covering metropolitan France using the SAFRAN, E-OBS, and ERA5-Land climate datasets. The results are presented as heat maps (Figure 5). The mean P50_id value obtained across the entire territory using the SAFRAN climate dataset was −3.37 MPa, with a wide Range of variation extending from −2.1 MPa in the wettest areas to −9.3 MPa along the Mediterranean region. P50_id was negatively correlated with climatic aridity indices, such as annual potential evapotranspiration (PET) (Figure 5, inset). However, grid points located in the Alpine region (blue dots) exhibited an opposite relationship with PET, highlighting the influence of extreme winter conditions on P50_id.

**Figure 5:**
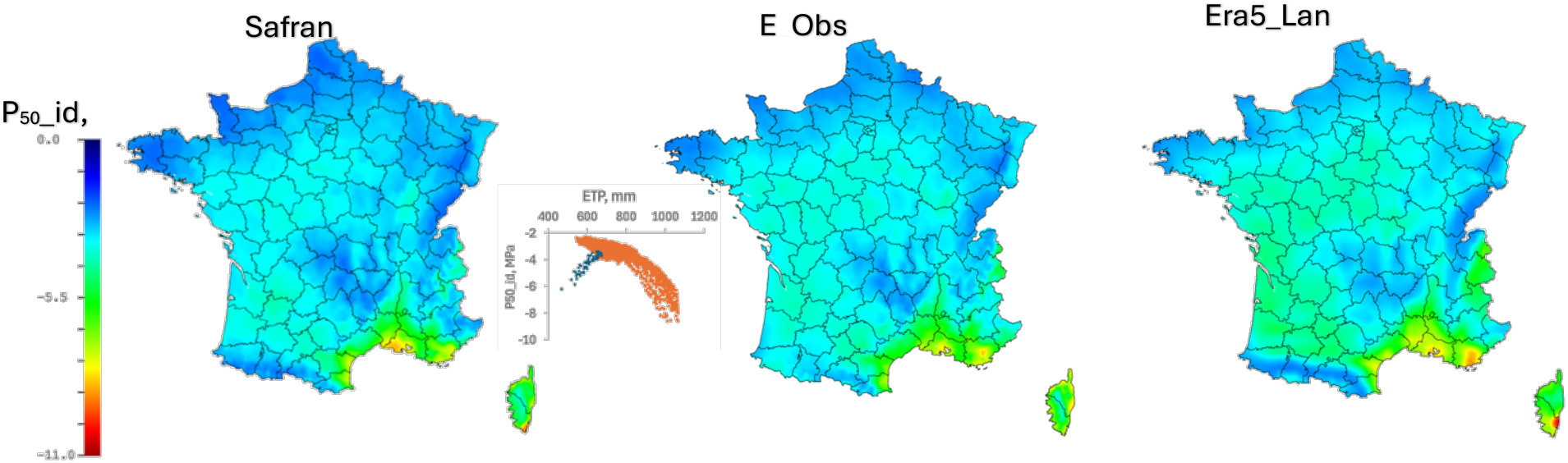
Spatial distribution of P_50__id across metropolitan France computed using SAFRAN, E-OBS, and ERA5-Land climate datasets. Results are shown as gridded maps illustrating strong spatial heterogeneity in ideotype xylem resistance to embolism. Insets show relationships between P_50__id and PET.

On average, P50_id values computed using the E-OBS climate dataset were highly correlated with those obtained using SAFRAN (figure 6), but were slightly more negative (−3.56 MPa). Differences were more pronounced when using ERA5-Land, which yielded even lower values (−3.79 MPa).

**Figure 6:**
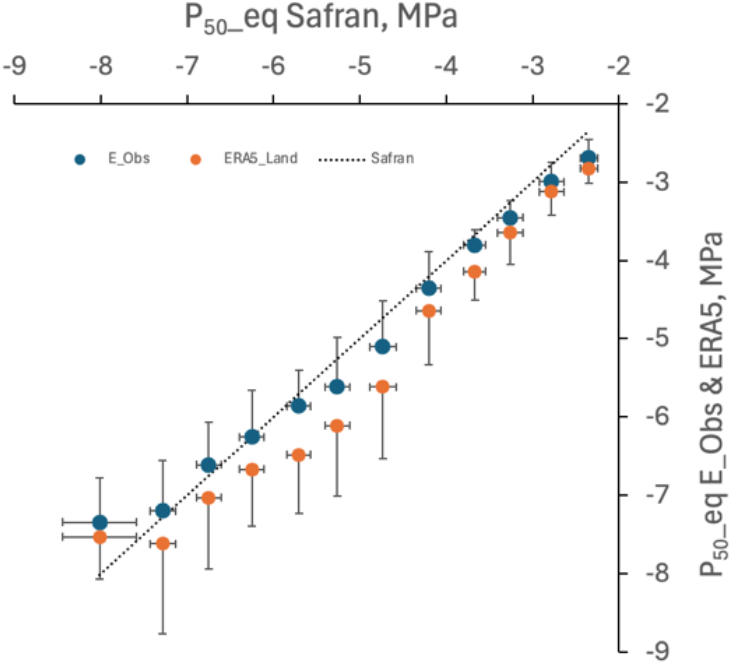
Comparison of spatially averaged P50_id estimates across metropolitan France obtained using different climate forcing datasets.

At the Euro–Mediterranean scale, the computation of P50_id reveals a strong latitudinal gradient, consistent with the climatic gradient and the distribution of major biomes. P50_id values are least negative in boreal and temperate biomes, and become increasingly negative in desert and savanna regions (figure 7). The two climate datasets tested (ERA5-Land and EC-Earth3) yield highly comparable results.

**Figure 7:**
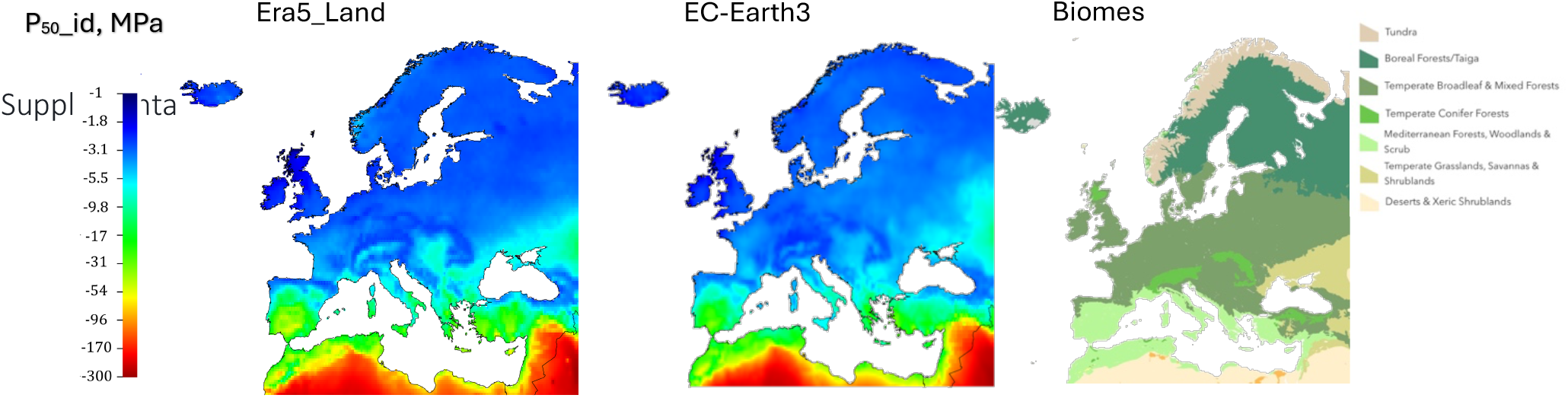
Spatial distribution of P50_id across the Euro–Mediterranean region derived from two climate datasets (ERA5-Land and EC-Earth3). The right-hand panel shows the distribution of major biomes across the region based on (c) RESOLVE Ecoregions, highlighting the correspondence between climatic gradients, biome structure, and ideotype xylem resistance to embolism.

### ISF esCmates and comparison with PDSI and SPEI drought indices

The ISF value was first calculated across France for the 1960–2025 period using the SAFRAN climate dataset. Mean ISF values over the country remained below 30% for most years (Figure 8), although several years exhibited exceptionally high ISF values, notably 1976, 2003, and 2022. As a consequence, the distribution of ISF values is strongly skewed around the mean (Figure 8, inset). The spatial distribution of ISF across France reveals strong geographic patterns that vary markedly among years. In 1976, exceptionally high ISF values were mainly confined to the northern half of the country, whereas in 2003 and 2022, high ISF values extended across nearly the entire territory (figure 8).

**Figure 8:**
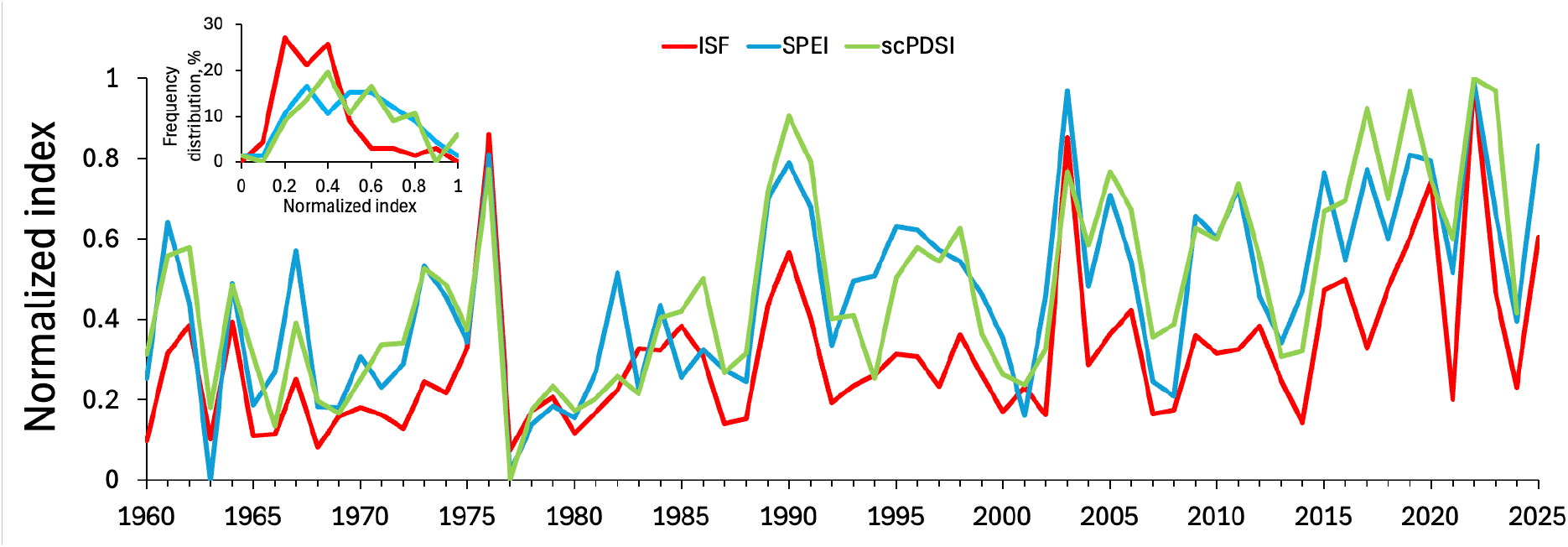
Interannual variability of ISF over metropolitan France (1960–2025) computed using the SAFRAN climate dataset. The inset shows the strongly right-skewed distribution of ISF values, reflecting the dominance of a limited number of extreme drought years. ISF is compared with normalized SPEI and scPDSI indices, both of which show significant correlations but more symmetric distributions across years.

**Figure 9:**
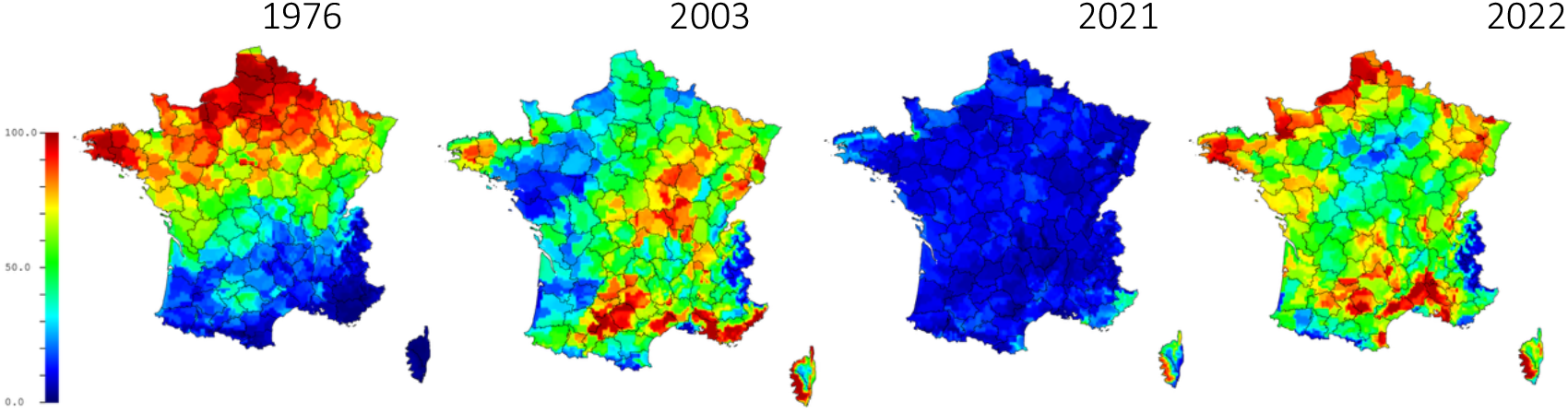
Spatial patterns of ISF across metropolitan France for selected extreme drought years (1976, 2003, and 2022) and a wet year (2021). The maps highlight strong interannual differences in the geographic extent and intensity of hydraulic stress, with localized stress in northern France in 1976 and near-countrywide high ISF values in 2003 and 2022.

We compared mean ISF values with two drought indices commonly used in forest ecology, SPEI and scPDSI. For comparison purposes, these indices were normalized between 0 and 1 in Figure 7. Both SPEI and scPDSI were significantly correlated with ISF, with R^2^ values of 0.66 and 0.62, respectively. However, the distributions of these indices were much more symmetric around their mean values, resulting in a larger number of years exhibiting relatively high drought index values.

At the Euro–Mediterranean scale, ISF also exhibited strong interannual regional variability (figure 10a). For example, high ISF values were mainly restricted to western Europe in 1976 and 2003, whereas they became much more widespread in 2018 and 2022. Overall, these spatial patterns remained broadly consistent with those observed for scPDSI (figure 10b) and SPEI (figure 10c). However, both drought indices tended to indicate high drought severity over more extensive areas than ISF.

**Figure 10:**
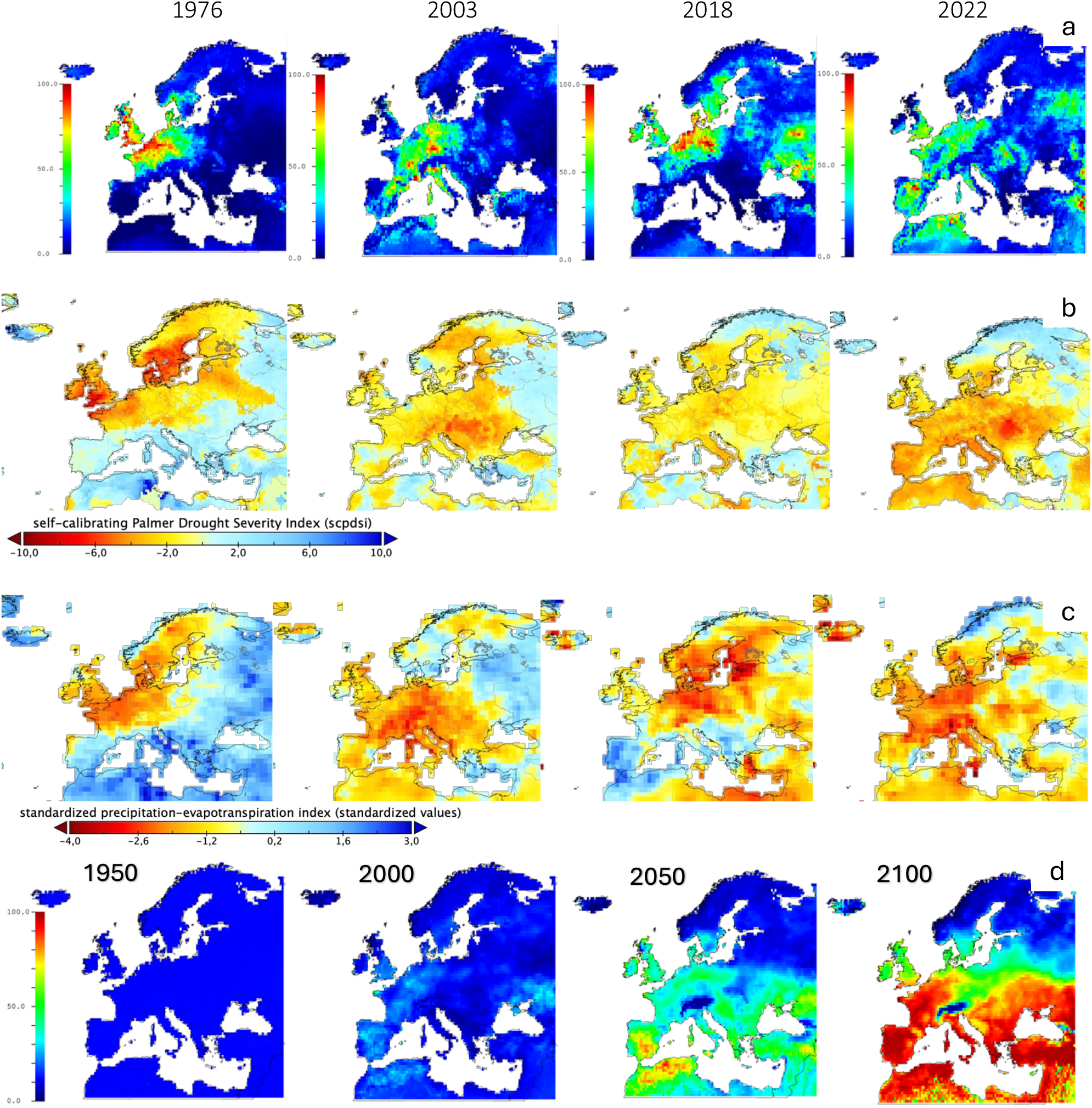
Spatial and temporal dynamics of ISF (a) scPDSI (b) and SPEI (c) across the Euro–Mediterranean region. (d) Projected evolution of ISF under the SSP5–8.5 scenario.

In a final analysis, we projected the evolution of ISF across the Euro–Mediterranean region up to 2100 under the SSP5–8.5 scenario (Figure 10d). Between 1950—the reference period during which ISF was, by construction, equal to 12%—and 2100, our simulations predict a progressive and dramatic increase in ISF across most of the territory, with the exception of the northernmost and alpine regions. In large parts of southern and central Europe, ISF reaches levels indicative of severe and recurrent hydraulic stress, suggesting that long-term tree survival under current trait syndromes would be increasingly compromised without substantial adaptation.

## Discussion

The ISF framework provides a mechanistic alternative to classical climatic drought indices by explicitly linking water stress to tree hydraulic functioning. Unlike SPI, PDSI or SPEI, which quantify atmospheric or soil water anomalies, ISF is grounded in the physiological response of trees to drought through xylem embolism dynamics. This constitutes a conceptual advance, as it directly connects climate forcing to a process closely associated with hydraulic failure and mortality risk.

The calculation of the ISF relies on the identification of an idealized tree phenotype in which the combination of key physiological traits minimizes the risk of embolism over a given reference period. We chose to base the determination of this ideotype on xylem resistance to embolism (P_50__id), while assuming empirical correlations between the other traits and P_50_. We showed that the value of P_50__id is inherently dependent on the parameterization of the SurEau model and, to a lesser extent, on the climatic datasets used. Thus, P_50__id strongly depends on leaf area index (LAI), tree size, soil water holding capacity, and the reference PLC threshold. Remarkably, however, ISF values are largely independent of these choices of SurEau model parameterization, making ISF a particularly robust and reproducible index.

The trait values we used to establish correlations with P50 are primarily representative of trees from temperate and semi-arid regions. In this context, the P50_id values obtained under our assumptions in the SurEau model parameterization are broadly consistent with those measured in species native to these regions. In contrast, the P50_id values simulated for Mediterranean and steppic biomes appear excessively negative. To our knowledge, the most negative P50 ever measured in a tree is approximately −18 MPa (Larter et al. 2015), and it has been suggested that more negative xylem pressures may be limited by the tensile strength of water molecules themselves (Caupin and Herbert 2006). This does not necessarily undermine the calculation of the ISF for these arid regions, but it likely indicates that other drought-resistance strategies than tolerance play a major role under such climates. These include drought escape strategies, access to deep water resources in phreatophytic species, and drought-deciduous behavior. Future developments of the ISF framework could therefore improve its applicability to arid regions by implementing alternative SurEau parameterizations that account for the specific drought-resistance mechanisms prevailing in these environments.

The agreement with SPEI and scPDSI further indicates that ISF captures the major climatic drought signals, while providing additional biological interpretation. However, a key difference lies in the statistical structure of the outputs : ISF exhibits a more skewed distribution and tends to emphasize extreme years rather than intermediate variability, suggesting a stronger sensitivity to climatic extremes and nonlinear physiological responses. This sensitivity to rare extreme events is particularly relevant in the context of our search for indicators of lethal water stress in forest biomes. Indeed, climatic episodes that have triggered severe forest dieback have been relatively infrequent since the 1950s (in France, for example : 1976, 2003, and 2022). It is therefore important to favor indices that are specifically sensitive to such rare but high-impact events. Under future climate scenarios (SSP5–8.5), ISF projects a marked and widespread increase in hydraulic stress across most of the Euro–Mediterranean domain, consistent with expectations of intensified atmospheric evaporative demand and precipitation deficits. Importantly, the model suggests a strong spatial divergence, with northern and alpine regions remaining relatively buffered, highlighting the role of climatic refugia under warming conditions.

A central strength of ISF lies in its process-based formulation. By embedding xylem embolism dynamics through the SurEau model, the index integrates key ecohydrological processes such as stomatal regulation, hydraulic conductivity loss, and soil–plant–atmosphere coupling. This allows ISF to go beyond “water availability” proxies and instead quantify stress in terms of functional disruption. Its main contribution lies in providing a mechanistically interpretable index of drought stress that is directly linked to a well-established pathway of tree mortality (Mantova et al 2023).

Future developments should focus on validating ISF values by establishing statistical relationships with empirical data on tree decline and mortality. This should help demonstrate the added value of the ISF compared to the climatic indices currently in use. Ultimately, ISF should be viewed not as a replacement for existing drought indices, but as a complementary tool that translates climatic anomalies into hydraulically meaningful stress indicators, thereby improving our ability to assess forest vulnerability under ongoing climate change.

## Acknowlegments

This work was supported in part by the International Research Center on Sustainable AgroEcosystems (ISITE CAP20-25, ANR-16-IDEX-0001), the TRONSAVE project (DSF/ONF), and the AURADAPT project (FAEDER PEI-Forêt).

---

**Figure s1:**
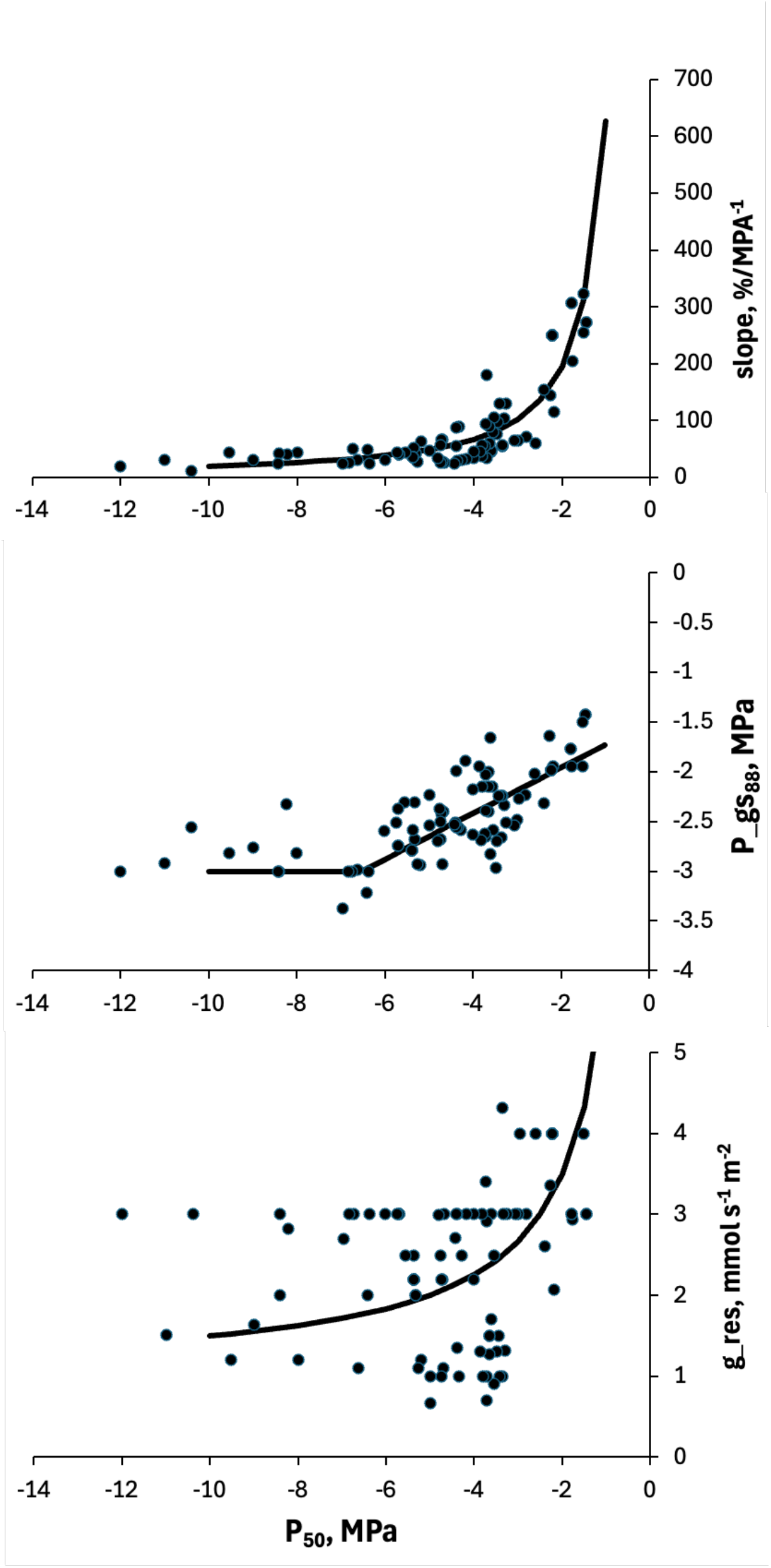
Empirical relationships between xylem vulnerability to em bolism (P_50_) and, the slope of the vulnerability curve at P_50_ (upper), the water potential inducing 88% stomatal closure (middle), and the residual leaf conductance (lower). Each point represents a different species. The curves show the regression used to parameterize these traits in the SurEau model for the calculation of P_50__id.

